# Alternative oxidase in trypanosomatids

**DOI:** 10.64898/2026.09.27.754813

**Authors:** Kristína Záhonová, Ondřej Gahura, Katarína Benediková, Natália Trusina, Elora Kalita, Ľubomíra Chmelová, Ingrid Škodová-Sveráková, Alena Zíková, Vyacheslav Yurchenko

## Abstract

**Background:** Alternative oxidase (AOX) is a mitochondrial terminal oxidase that provides an alternative route for electron transfer, contributing to respiratory flexibility and redox homeostasis under stress conditions. In trypanosomatids, AOX displays a highly uneven distribution associated with extensive diversification of mitochondrial electron transport chains (ETCs). In addition to canonical AOX, many trypanosomatids encode an enigmatic AOX-like protein (AOX-L), whose evolutionary origin and functional significance remain unresolved.

**Results:** Here, we investigated the evolutionary distribution, phylogenetic relationships, structural conservation, and functional divergence of AOX and AOX-L across euglenozoans. Comparative genomic analyses revealed a patchy distribution of both proteins among kinetoplastids, with multiple independent losses during evolution. Phylogenetic analyses demonstrated that AOX and AOX-L represent distinct evolutionary lineages, indicating independent origins rather than duplication-derived divergence. While AOX showed a conserved eukaryotic origin, AOX-L occupied a separate phylogenetic position and lacked key residues required for ubiquinol oxidation. Structural modelling revealed that AOX-L retained the characteristic AOX fold, predicted membrane association, and dimeric organization, but lacked the catalytic architecture necessary for enzymatic activity. Transcriptomic and biochemical analyses of four AOX-containing trypanosomatid species showed that AOX expression and activity correlated with mitochondrial ETC organization. Species lacking cytochrome-dependent complexes III and IV displayed substantially higher AOX expression and enzymatic activity, consistent with AOX functioning as the primary terminal oxidase in these lineages. Conversely, species retaining a canonical ETC exhibited lower AOX activity, suggesting a role in metabolic flexibility and redox regulation.

**Conclusions:** Our study provides a comprehensive evolutionary and functional framework for alternative oxidases in trypanosomatids. We demonstrate that AOX and AOX-L are evolutionarily distinct proteins with different predicted functions: AOX maintains respiratory electron flow according to lineage-specific mitochondrial requirements, whereas AOX-L represents a structurally conserved but catalytically inactive protein family that may have acquired an alternative regulatory role. These findings highlight how mitochondrial respiratory components diversify during eukaryotic evolution and provide a basis for future investigations into the biological function of AOX-L.

## Background

Alternative oxidase (AOX) is a mitochondrial salicylhydroxamic acid (SHAM)-sensitive oxidoreductase. It catalyzes reduction of oxygen to water by accepting electrons from the ubiquinol pool in the electron transport chain (ETC) [1]. Unlike the cyanide-sensitive cytochrome *c* oxidase (COX, complex IV), AOX does not translocate protons across the inner mitochondrial membrane and, therefore, does not contribute to ATP generation by oxidative phosphorylation. Instead, it provides an alternative pathway for electron transfer, bypassing the complexes III and IV of canonical ETC. Although this reduces the efficiency of ATP production, it maintains electron flow, limits the formation of reactive oxygen species (ROS) and helps preserving cellular redox balance under conditions of respiratory stress or COX pathway inhibition [2-4]. This, in turn, implicates AOX in different cellular processes, such as redox homeostasis, heat and oxidative stress responses, and metabolic adaptation across diverse eukaryotes [1, 5, 6].

Alternative oxidase has a remarkably patchy distribution across the tree of life. Whereas it is restricted to proteobacteria within prokaryotes, it has a highly variable occurrence within eukaryotes, with numerous independent losses and instances of horizontal gene transfer, indicating that its function is beneficial only under specific physiological conditions [7, 8]. This evolutionary divergence is also reflected in the biochemical properties of the enzymes that can be modulated by, for example, oxoacids and purine nucleotides.

Structurally, alternative oxidase is a homodimeric protein that is peripherally associated with the matrix-facing leaflet of the inner mitochondrial membrane [9, 10]. Each monomer of AOX consists of six α-helices forming a conserved α-helix bundle with a buried active site and two hydrophobic cavities to bind its substrate, ubiquinol [11]. The di-iron catalytic center is coordinated by four glutamate (E) and two histidine (H) residues, and is characterized by two conserved ExxH motifs, typical of the di-iron metalloproteins [10, 12, 13]. The AOX proteins generally have molecular weights ranging between 32-38 kDa and maintain their highly conserved catalytic core despite sequence divergence across taxa [1].

Notably, the fungi-like AOX has also been documented in several parasitic protists of the family Trypanosomatidae, where its activity is regulated by mitochondrial ATP levels [14-17]. Trypanosomatids (Euglenozoa: Kinetoplastea: Trypanosomatidae) constitute a group of flagellated, obligatory parasitic protists that infect different organisms, including plants, invertebrates, and vertebrates [18, 19]. Some of them are medically relevant and clinically significant as they cause serious, but often neglected, human diseases, such as sleeping sickness (*Trypanosoma brucei*), Chagas disease (*T. cruzi*), and leishmaniasis (*Leishmania* spp.) [20-22]. They are non-taxonomically divided into monoxenous (with one host in their life cycle) and dixenous (with two hosts in their life cycle: genera *Leishmania sensu lato*, *Phytomonas*, and *Trypanosoma*) species [23, 24]. Adaptation to different host environments and ecological niches has driven the diversity of trypanosomatids’ gene repertoire and, consequently, their metabolic capacities [25]. This is well reflected in the composition and activity of the mitochondrial ETC complexes across trypanosomatid species and developmental stages [26-30]. The AOX in trypanosomatids was traditionally referred to as trypanosomatid alternative oxidase (TAO) [16], however, to keep consistency with other works, we use AOX hereafter. Even within this clade, AOX has a patchy distribution that closely mirrors the diversity of their ETC. For example, *Leishmania* spp. possess a canonical cytochrome-dependent ETC with complexes III and IV and no AOX [14, 31, 32]. *Trypanosoma brucei* encodes both AOX and a canonical cytochrome-dependent ETC complexes III and IV. The developmental regulation of these two respiratory systems enables this organism to adapt to the metabolically diverse environments of the tsetse fly and the mammalian host [33, 34]. The insect-stage procyclic form (PCF) relies predominantly on complexes III and IV for mitochondrial respiration and expresses only low levels of AOX. Conversely, the mammalian bloodstream form (BSF) lacks cytochrome-containing complexes III and IV, making AOX the sole terminal oxidase of its truncated ETC [16, 35, 36]. Similarly to *T. brucei* BSF, dixenous *Phytomonas* spp. [14] and monoxenous *Vickermania* spp. [37] also lack cytochrome-containing ETC and rely on AOX for terminal electron transfer [38, 39]. The presence of AOX in several pathogenic protists and fungi, but not in mammalian cells, makes this enzyme a promising potential drug target [40-42].

Analysis of the functional importance of AOX in trypanosomatids is further complicated by the presence of another, arguably even more enigmatic, AOX-like protein, previously termed AOX2 or TAO-like, encoded in genomes of many species [16]. Although AOX-L shares sequence similarity with AOX, previous analyses have suggested that it lacks several residues required for catalysis, raising the possibility that it has evolved a distinct, non-enzymatic function. Yet, its evolutionary origin, structural conservation, and relationship to canonical AOX remain unresolved. Here, we investigated phylogenetic distribution of AOX and AOX-L across trypanosomatids and used a structural modelling approach to assess their evolutionary conservation and functional divergence. Our findings provide a framework for understanding the emergence and diversification of alternative oxidases and their adaptation to the diverse mitochondrial respiratory strategies across trypanosomatid lineages.

## Methods

### Sequence analyses, phylogenetic inference, and structural modelling

AOX and AOX-L sequences were identified by BLAST v. 2.9.0+ [43] searches integrated in the AMOEBAE workflow [44] using *T. brucei* proteins as queries and genomes, genome-derived proteomes, and transcriptomes of euglenozoans from NCBI, TriTrypDB release 68 [45], MMETSP [46] (reassembly available at https://doi.org/10.5281/zenodo.257410), and previous publications [47-51] as databases (Table S1). Identified transcript sequences in the diplonemid *Paradiplonema papillatum* were confirmed in the high-quality genome assembly [52]. Note that two identified transcripts, Dpap-TRINITY_DN39793_c0_g8389_i5 and Dpap-TRINITY_DN12048_c0_g3740_i3 correspond to the genomic loci KAJ9448012 and KAJ9471081, respectively. Two additional genome-derived proteins (KAJ9448010 and KAJ9448011) were found to be fragmentary and, therefore, excluded from further analyses. Subcellular localizations were predicted by TargetP v. 2.0 [53] and DeepLoc v. 2.1 [54].

Identified euglenozoan sequences were added to the AOX and terminal oxidases datasets from previous works [7, 55]. All sequences were aligned by MAFFT v. 7.310 [56] under L-INS-i method and trimmed with trimAl v. 1.4.rev15 [57] using -gt 0.8 option to remove poorly aligned and ambiguous positions. Maximum-likelihood phylogenetic analysis was conducted in IQ-TREE v. 2.2.0 [58] with LG+G4 model for the guide tree, followed by the posterior mean site frequency analysis [59] using LG+C20+G4 model and the guide tree input with 1,000 replicates for ultrafast bootstraps [60] and a maximum of 5,000 iterations.

Structures of AOX and AOX-L monomers and dimers were predicted by AlphaFold3 [61] and superposed, visualized, and analyzed in UCSF ChimeraX [62].

### Transcriptomic analysis

For evaluating expression levels of AOX and AOX-L, the TPM (transcripts per million) values were calculated. For *T. brucei* and *A. deanei*, available protein-coding sequences from the TriTrypDB were used. For *V. ingenoplastis*, protein-coding sequences published previously [37] were used. For *P. françai*, protein-coding genes were predicted by Braker v. 3.0.8 [63] using the published genome assembly [64] and a database of kinetoplastid proteins enriched with sequences of *Blastocrithidia nonstop* and *Obscuromonas modryi* [65, 66].

The TPM values were calculated from the RPKM (reads per kilobase of transcript per million reads mapped) values using the following formula: TPM = (RPKM of a transcript)/(sum of RPKMs of all transcripts) × 10^6^ (Table S2). Firstly, RNA-seq reads (SRA accessions: SRR1272139, SRR1272140, and SRR1272141 for *T. brucei* PCF; SRR1986388, SRR1986389, and SRR1986390 for *A. deanei*; ERR1655128 and ERR1655129 for *P. françai*; SRR31549352 for *V. ingenoplastis*) were adapter and quality trimmed by BBDuk v. 39.01 (part of BBTools suite [67]) and mapped onto the protein-coding sequences using BBMap v. 39.01 (part of BBTools suite). The RPKM values were obtained directly from BBMap using “rpkm= <FILE<” parameter. If several genomic loci encoded the protein, their TPM values were summed.

### Cultivation of trypanosomatid species

Procyclic form (PCF) of *T. brucei* strain Lister 427 line 29-13 was grown at 27 °C in SDM-79 (Thermo Fisher Scientific, Waltham, USA), supplemented with 10% (v/v) heat-inactivated fetal bovine serum (Biosera, Cholet, France), 2 μg/ml hemin (Merck, Rahway, USA), 50 U/ml penicillin, and 50 μg/ml streptomycin (both from Jena Bioscience, Jena, Germany). *Vickermania ingenoplastis* strain CP21, *Angomonas deanei* strain CT-IOC-044, and *Phytomonas serpens* strain 9T were grown at 23 °C in Schneider’s *Drosophila* medium (Biosera, Cholet, France) supplemented as above. Species identity was validated using 18S rRNA locus sequencing as described previously [68].

### Mitochondrial fraction isolation and AOX activity assay

Cells were pelleted (5,000× *g*, 10 min, 4 °C) at the end of their exponential phase of growth, washed in ice-cold STE buffer (250 mM sucrose, 20 mM Tris-HCl pH 7.9, and 2 mM EDTA), aliquoted, and stored at -80 °C. Mitochondria from each aliquot (5× 10^8^ cells) were isolated by differential centrifugation from cells lysed using hypotonic lysis as described previously [69]. Isolated mitochondria were resuspended in 0.5 M aminocaproic acid and 2% (w/v) dodecylmaltoside and kept on ice for 1 h. The lysate was then centrifuged at 25,000× *g* for 10 min at 4 °C, and the protein concentration was determined by Bradford method [70].

Before starting the measurement, mitochondrial lysates were incubated in the assay buffer (50 mM Tris-HCl pH 7.4) in a 1 ml silica cuvette for 2 min, 10 µl of 10 mM reduced Q_2_ was added, and the change in ubiquinone absorbance was measured for 2.5 min at 278 nm every 10 s at 25 °C [71]. Activity unit (U) is defined as the amount of enzyme that utilizes 1 μmol of substrate per minute. The given values are presented as an average of 3-4 independent biological replicates, each containing at least 3 technical replicates calculated per 1 mg of mitochondrial proteins. The specificity of AOX activity was confirmed in the presence of its inhibitor (0.1 mM SHAM). In mitochondrial lysates of *T. brucei*, over 90% of activity was SHAM-dependent. In mitochondrial lysates of *A. deanei*, the measured AOX activity was less (only about 30%) sensitive to SHAM. The remaining activity, thus, may belong to complex III (ubiquinol-cytochrome *c* oxidoreductase), which competes for the same substrate. Complex III is not present in *P. serpens* and *V. ingenoplastis*; and, as such, AOX is the only enzyme with ubiquinole oxidase activity present in their mitochondrial lysates.

## Results

### Presence of AOX and AOX-L in euglenozoans

Firstly, we sought to determine whether both AOX and AOX-L genes in kinetoplastids are of eukaryotic origin. We queried 48 kinetoplastid, ten diplonemid, and three euglenid datasets for the presence of these genes (Fig. 1; Table S1). While diplonemids and euglenids uniformly encode AOX (in some cases, in several copies), none of these organisms possess AOX-L. On the other hand, the distribution of AOX and AOX-L in kinetoplastids is patchy. The majority of investigated organisms have only AOX-L, four (*Vickermania ingenoplastis* and three *Phytomonas* spp.) encode only AOX, five (*Angomonas deanei*, *Bodo saltans*, *Trypanosoma brucei*, *T. congolense*, and *T. vivax*) possess both AOX and AOX-L, and only two (*Blastocrithidia nonstop* and its close phylogenetic relative *Obscuromonas modryi*) do not encode either. The observed distribution implies at least nine and three independent losses of AOX and AOX-L, respectively, in the evolutionary history of kinetoplastids (Fig. 1).

**Fig. 1.**
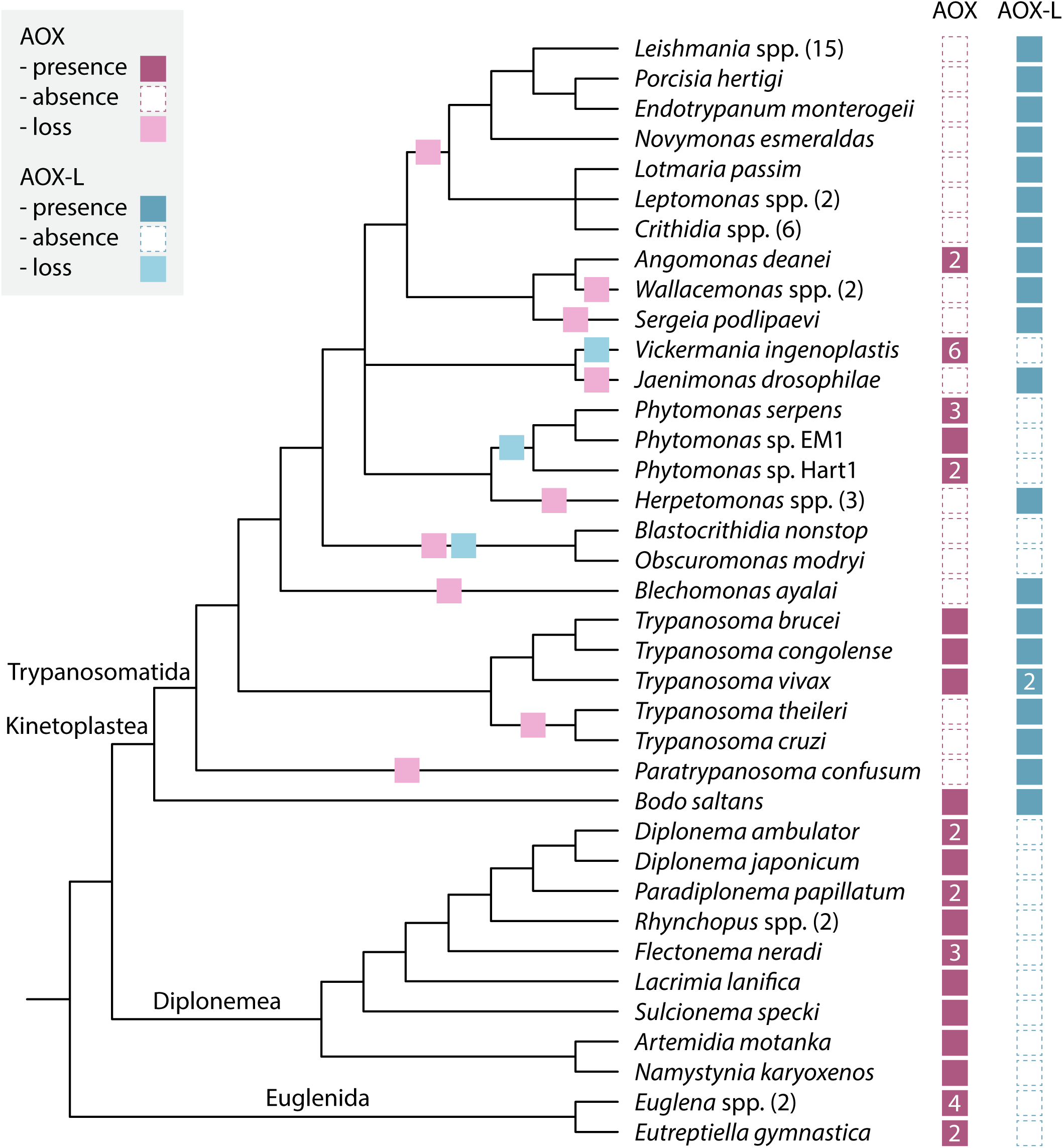
Distribution of AOX and AOX-L in euglenozoans. Identified and missing proteins are shown as colored and white squares, as explained in the graphical legend. Numbers in parentheses indicate number of species of the same genus examined, whereas numbers in squares indicate number of identified sequences (Table S1). Gene losses are mapped onto a schematic phylogenetic tree based on previous works [18, 25].

We next performed phylogenetic analysis (Fig. S1; Data S1) taking advantage of previously published comprehensive datasets [7, 55]. The included plastid terminal oxidases served as an outgroup and the rooting point. As in the previous analysis [7], bacterial sequences formed a monophyletic clade inside eukaryotic AOX. The position of three euglenid clades was consistent with the previous work [55] corroborating robustness of the tree. Diplonemid sequences formed two clades close to the majority of euglenid AOX. Kinetoplastid AOX sequences were positioned close to those of stramenopiles. Kinetoplastid AOX-L sequences were previously nested inside the bacterial clade [7], however, now they formed a basal clade to all eukaryotic and bacterial AOX hindering further affiliation.

In summary, we have identified four trypanosomatid lineages encoding AOX enzymes in their genomes – *Angomonas*, *Phytomonas*, *Trypanosoma*, and *Vickermania*. Please note that in trypanosomes, AOX presence appears to be restricted to subgenera *Trypanozoon* and *Duttonella*, although many other *Trypanosoma* spp. were not analyzed because they lack genomic data. Our results strongly suggest that kinetoplastid AOX- and AOX-L-encoding genes are not phylogenetically related and emerged independently: *Aox* is of eukaryotic origin, while phylogenetic position of *Aox-l* is basal to all analyzed eukaryotes and bacteria.

### Analysis of AOX expression and enzymatic activity

Given the variable organization of the mitochondrial respiratory chain in trypanosomatids [30, 37, 39, 72-74], we chose four AOX-possessing species differing in the presence/absence of AOX-L to monitor expression levels of both genes and enzymatic activity of AOX (as mentioned above, the AOX-L protein is enzymatically inactive). *Trypanosoma brucei* and *A. deanei* encode both proteins, whereas *P. serpens* and *V. ingenoplastis* possess only AOX (Fig. 1). For evaluating the AOX and AOX-L expression, we calculated TPM values (Fig. 2A; Table S2). Since no transcriptomic data are publicly available for *P. serpens*, we used data for a phylogenetically closely related *P. françai*. Both procyclic *T. brucei* and *A. deanei* exhibited low levels of expression of AOX (17 and 23 TPM, respectively) as well as AOX-L (20 and 31 TPM, respectively). On the other hand, the AOX expression was much higher in *P. françai* and *V. ingenoplastis* (890 and 1,630 TPM, respectively).

**Fig. 2.**
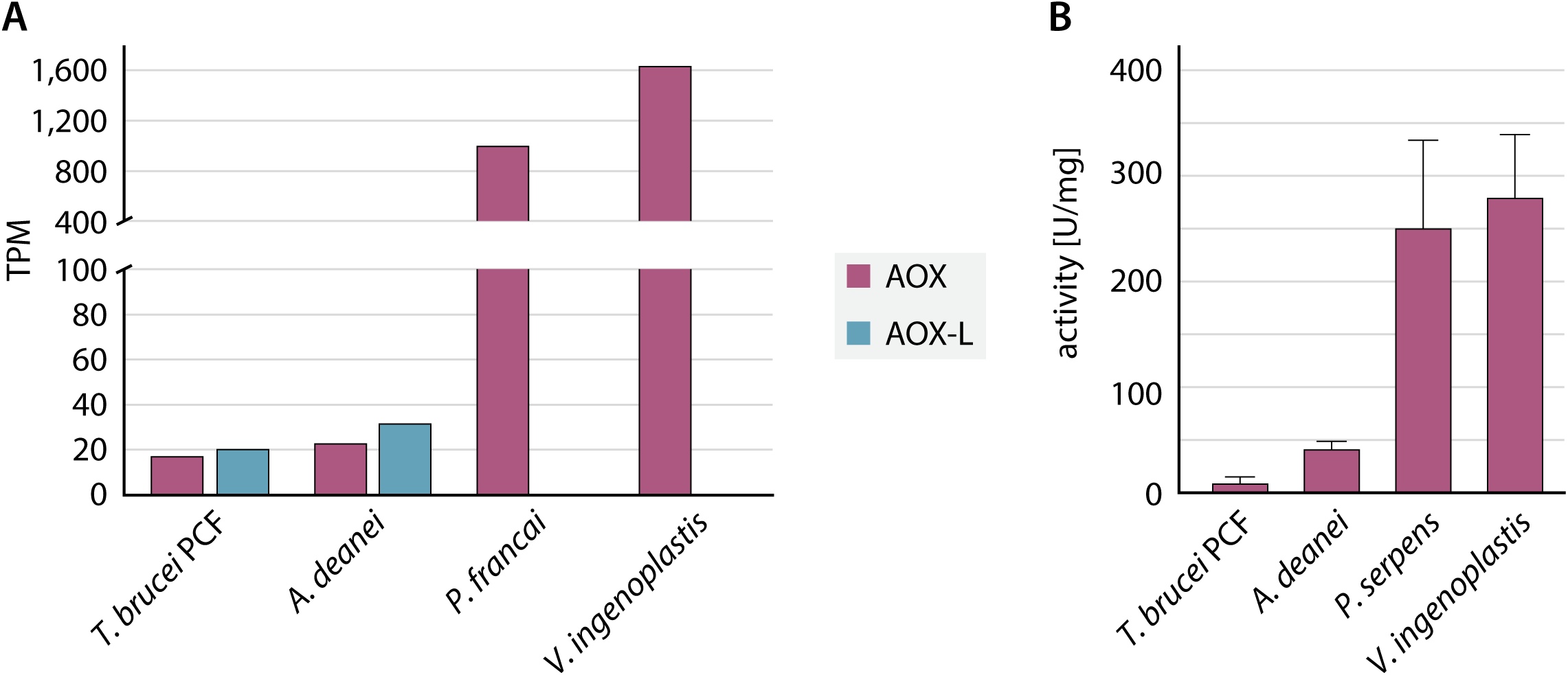
AOX expression and activity levels in selected trypanosomatids. **(A)** The expression of AOX and AOX-L, showed as TPM values, were calculated from RPKM using the formula: TPM = (RPKM of a transcript)/(sum of RPKMs of all transcripts)× 10^6^ (Table S2). **(B)** Enzymatic activity of AOX in mitochondrial lysates of *T. brucei* procyclic form (PCF), *A. deanei*, *P. serpens*, and *V. ingenoplastis.* Specific activity is expressed as μmol of substrate converted per minute per mg of mitochondrial protein (mU/mg). Given values represent an average of at least three independent measurements, each containing at least three technical replicates.

To assess the enzymatic ubiquinol oxidase activity in four species possessing AOX, Q_2_ oxidation in mitochondrial lysates of *A. deanei*, *P. serpens*, procyclic *T. brucei*, and *V. ingenoplastis* was analyzed spectrophotometrically (Fig. 2B). In concert with the transcriptomic data, *V. ingenoplastis* and *P. serpens* had the highest specific activity (282 and 253 mU/mg, respectively). The activity was approximately six-fold lower in *A. deanei* (44 mU/mg), and the lowest in the *T. brucei* (12 mU/mg).

### Detailed analyses of sequences and predicted structures of AOX and AOX-L proteins

Finally, to assess whether the observed differences in AOX enzymatic activities could be explained by sequence and/or structural divergence, we compared sequences and available experimental or AlphaFold-predicted structures of AOX from *A. deanei*, *P. serpens*, *T. brucei*, and *V. ingenoplastis*. On the sequence level, we identified five regions with relatively high divergence (Fig. S2). The first one is located within the N-terminal arm, in a region connecting two short α-helices contributing to the dimer interface in the *T. brucei* crystal structure [10]. In *Phytomonas* spp.*, A. deanei*, and *V. ingenoplastis*, this region contains an insertion of four residues, resulting in a slight extension of the two short α-helices and overall local structure expansion and deviation. Three additional variable regions map to helices α5 and α6, however, these lie outside the di-iron carboxylate site or ubiquinone binding pocket, and, therefore, are unlikely to directly affect catalysis (Fig. 3). Finally, a C-terminal extension, specific to *T. brucei* and *T. congolense*, is unresolved in the crystal structure and unreliably predicted by the AlphaFold, suggesting that it is intrinsically flexible or disordered.

**Fig. 3.**
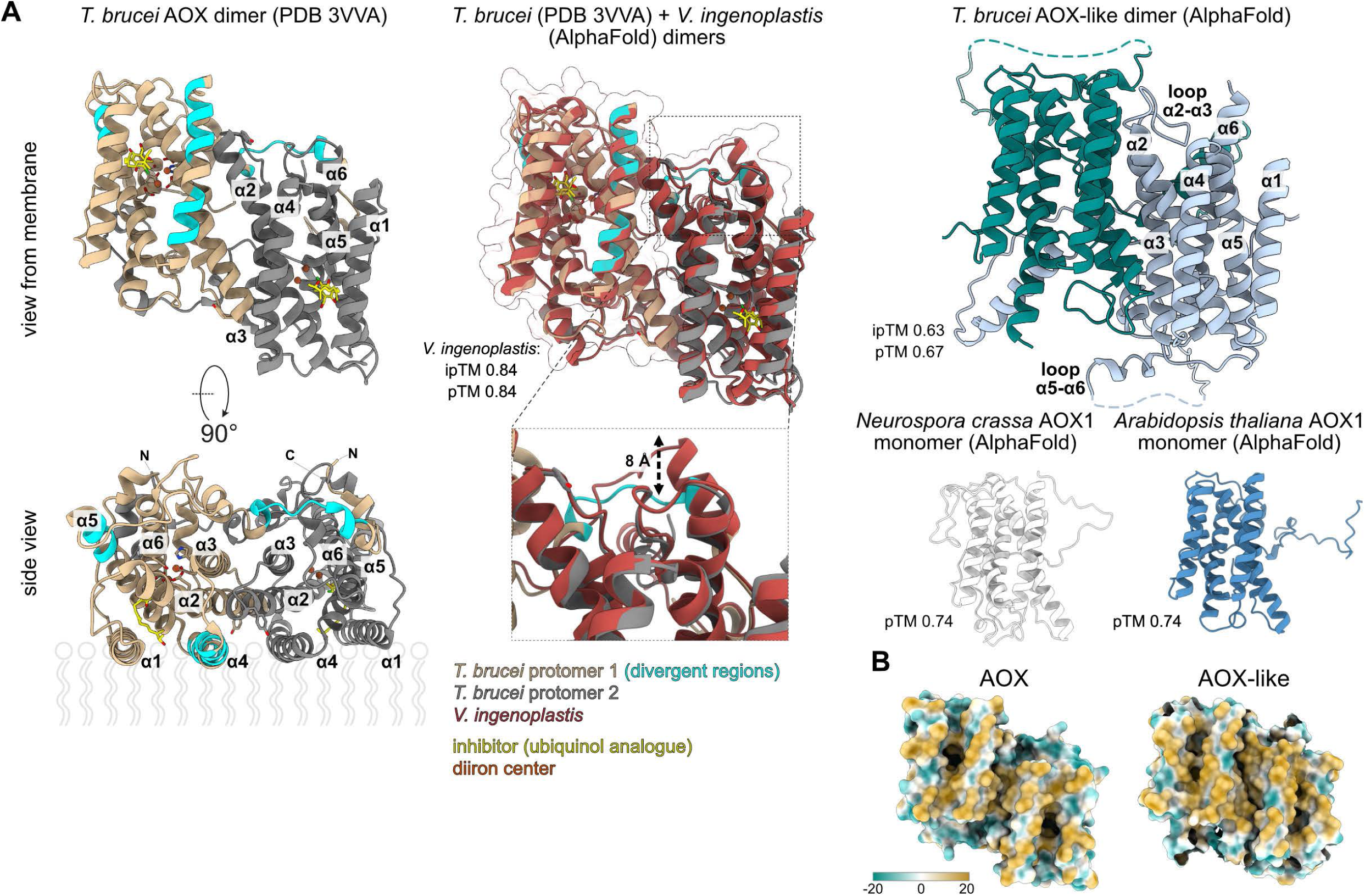
Comparison of structures of AOX and AOX-L proteins from selected trypanosomatids. **(A)** Structure of *T. brucei* AOX dimer with ubiquinol analog resolved by X-ray crystallography (PDB ID 3VVA) compared to AlphaFold-predicted models of dimers or monomers of AOX and AOX-L proteins from indicated organisms. Membrane is indicated in the side view. For all AlphaFold models, pTM scores are shown, and ipTM scores are shown for dimers. **(B)** View of *T. brucei* AOX and AOX-L from the membrane showing surface colored by molecular lipophilicity potential. Highly lipophilic regions are shown in dark golden rod and hydrophobic regions in cyan.

To further investigate the structural divergency between AOX and AOX-L, we compared AlphaFold-predicted models of both proteins from representative trypanosomatids with the experimentally determined structure of *T. brucei* AOX [10], as well as with predicted structures of AOX1 from the plant *Arabidopsis thaliana* (Uniprot ID Q39219), and the filamentous fungus *Neurospora crassa* (Q01355) (Fig. 3A). Despite their considerable sequence divergence, all AOX and AOX-L proteins share the characteristic AOX fold consisting of six antiparallel α-helices (α1-α6) with an N-terminal arm containing two to three short α-helices, and two additional short α-helices flanking α6. Thus, the overall structural framework of AOX has been largely retained in AOX-L.

The AOX-L proteins contain extended loops without predicted structures between helices α2 and α3, and α5 and α6. While the latter is common to all kinetoplastids, the former emerged most likely in the common ancestor of the branches containing *Phytomonas*, *Vickermania*, and *Leishmania*. Hydrophobic surfaces of helices α1 and α4 that presumably allow interaction of AOX with the inner mitochondrial membrane [9, 10] are also predicted in the AOX-L structure (Fig. 3B), implying that it binds the membrane in the same manner. In line with this, the AOX-L, similarly to the AOX, is highly enriched in the integral membrane fraction, as revealed by the recent proteomic study [75]. In contrast to this overall structural conservation, the AOX-L protein lacks five (E123, E162, E213, Y220, and H269 in *T. brucei* numbering) out of seven di-iron carboxylate active site residues conserved in AOX that are critical for the enzymatic activity. About half of the residues implicated in ubiquinone binding [10] are also absent in all AOX-L proteins. In addition, the predicted entry site to the substrate cavity appears to be constricted in AOX-L, suggesting that ubiquinone binding by AOX-L is rather unlikely.

The canonical AOX proteins in other organisms homodimerize [76-78]. As judged from the high interface predicted template modeling (ipTM) values (ranging between 0.63 and 0.84) of all AlphaFold dimer models (Fig. 3A), all trypanosomatid AOX and AOX-L presumably also occur as dimers. The inter-protomers interfaces are formed mostly by helices α2 and α3 and are further contributed by α4 and elements of the N-terminal arm. However, specific residues involved in dimerization are very different in the AOX and AOX-L molecules. In line with this observation, ipTM values of AlphaFold-predicted AOX/AOX-L heterodimers in *T. brucei* and *A. deanei* are only 0.12 and 0.13, respectively. Therefore, the existence of the heterodimers in the organisms, where both proteins co-occur, is unlikely.

Taken together, despite several localized sequence insertions and variable regions, the overall fold of AOX, including the catalytic core and substrate-binding architecture, is highly conserved across all four analyzed trypanosomatid species. Thus, the observed differences in enzymatic activity are unlikely to arise from major structural differences in the catalytic domain. Our analyses indicate that AOX-L has retained the overall architecture, membrane association, and oligomeric organization of the canonical AOX, while losing the structural determinants required for ubiquinol oxidation. This supports the view that AOX-L represents a structurally conserved but functionally divergent member of the AOX protein family.

## Discussion

The evolutionary history of trypanosomatids, as of other eukaryotes, has been shaped by gene duplications, acquisitions, and extensive gene loss events [79]. Transition from their non-parasitic free-living ancestor to parasitic lifestyle by these organisms has been accompanied by substantial genome reduction [23, 80]. The evolutionary consequences of this reductive selection are evident in the remarkable diversity of mitochondrial metabolism across trypanosomatids, including the uneven distribution of alternative oxidase. In this study, we focused on *Aox* and *Aox-l* genes, which were independently lost several times in the evolution of Trypanosomatidae. It may appear that AOX-L represents a nonfunctional or pseudogenized copy of AOX because many residues of the di-iron carboxylate active site implicated in ubiquinone binding are missing in the AOX-L sequences.

However, our phylogenetic analysis clearly showed that AOX and AOX-L are phylogenetically unrelated, meaning that they emerged independently in the evolution of trypanosomatids. In line with this and as judged from the AlphaFold modelling, while both AOX and AOX-L proteins are predicted to dimerize, the existence of AOX-AOX-L heterodimers is rather unlikely.

Notably, AOX enzymatic activity was substantially lower in *Angomonas* and *Trypanosoma*, both of which possess a canonical cytochrome-dependent ETC, than in *Phytomonas* and *Vickermania*, where complexes III and IV have been independently lost and AOX functions as the sole terminal oxidase. The observed differences closely parallel AOX transcript abundance, suggesting that AOX expression has been tuned to the bioenergetic requirements imposed by the organization of the ETC rather than by intrinsic differences in enzyme structure. Although the two species with lower AOX activity also encode AOX-L, our data do not imply a role for AOX-L in regulating AOX activity. Instead, the correlation is more readily explained by differences in the ETC architecture. Nevertheless, the evolutionary conservation of AOX-L, despite the apparent loss of its catalytic function, suggests that it has acquired an alternative, yet unidentified, role.

A scenario, in which a nonfunctional copy modulates the activity of an enzyme, is not unprecedented [81, 82]. This also concerns trypanosomatids. In *T. brucei*, for example, the activity of *S*-adenosylmethionine decarboxylase (AdoMetDC) is increased upon dimerization with AdoMetDC pseudoenzyme, called prozyme [83]. On post-transcriptional level, pseudogene-derived small interfering RNAs were shown to modulate gene expression *via* RNA interference [84]. We may have uncovered another example of such regulation with the *Aox* and *Aox-l* gene pair. The transcription in trypanosomatids is polycistronic with large clusters of functionally unrelated genes transcribed at once into a single pre-mRNA. The pre-mRNA is then split into individual mature mRNAs by *trans*-splicing [85, 86]. The stability of mRNAs is controlled by RNA-binding proteins, many of which bind to the 3′ UTRs enriched in low-complexity sequences, promoting protein translation [87-90]. Changes in any of these processes may also explain the observed differences in expression of AOX and AOX-L in the studied species.

Yet, the most parsimonious explanation of the documented phenotypes stems from the parasite biochemistry. In *Phytomonas* and *Vickermania*, complexes III and IV were independently lost leaving AOX as the only terminal oxidase [37, 38]. Consistent with this reorganization of ETC, a high AOX activity is expected to accommodate the electron flow redirected to AOX. The functional role of AOX in *Angomonas*, which maintains a canonical mitochondrial respiratory chain [91], remains an open question and warrants further investigation. It may represent a flexible regulatory element enabling the modulation of the mitochondrial redox state or protection against an excess of reducing equivalents and the formation of ROS, similarly to the situation observed in algae and plants [5, 92]. It might as well be involved in the control of bacterial endosymbionts in these species [93, 94].

Collectively, our analyses provide a revised view of alternative oxidases in trypanosomatids. The AOX and AOX-L represent evolutionarily distinct protein families that differ in both their origin and predicted function. The distribution and activity of AOX closely reflect the organization of the mitochondrial ETC. In lineages that have lost complexes III and IV, AOX functions as the sole terminal oxidase and exhibits high expression and activity to sustain respiratory electron flow. Conversely, in species retaining a canonical electron transport chain, AOX is maintained at lower abundance, where it likely acts as an alternative electron sink that enhances metabolic flexibility and contributes to maintaining redox homeostasis under changing physiological conditions. In contrast, AOX-L has retained the characteristic structural fold of AOX while losing the catalytic features required for ubiquinol oxidation, suggesting that it has evolved toward a distinct non-enzymatic function. Together, these findings establish a framework for understanding how mitochondrial respiratory proteins have diversified during the evolution of euglenozoans and provide a basis for future studies aimed at uncovering the biological role of AOX-L.

## Data availability

Predicted proteins of *Phytomonas françai* were deposited in Figshare: https://doi.org/10.6084/m9.figshare.32626794.

## Ethics, Consent to Participate, and Consent to Publish declarations

Not applicable.

## Acknowledgements

This work was primarily supported by the EU’s Operational Program ‘Just Transition’ (CZ.10.03.01/00/22_003/0000003 LERCO) to VY and the Ministry of Education, Research, Development and Youth of the Slovak Republic (VEGA 1/0709/26) to IS-S. Additional funding was provided by the EU’s Operational Program ‘Jan Amos Komenský’ (CZ.02.01.01/00/22_008/0004575 RNA for therapy) to AZ and OG, and Horizon-Europe ERC (101044951 MitoSignal) to AZ. Computational resources were provided by the e-INFRA CZ project (ID: 90254), supported by the Ministry of Education, Youth and Sports of the Czech Republic.

