## Supplementary figures and images for "Alternative oxidase in trypanosomatids"

### Fig. S1

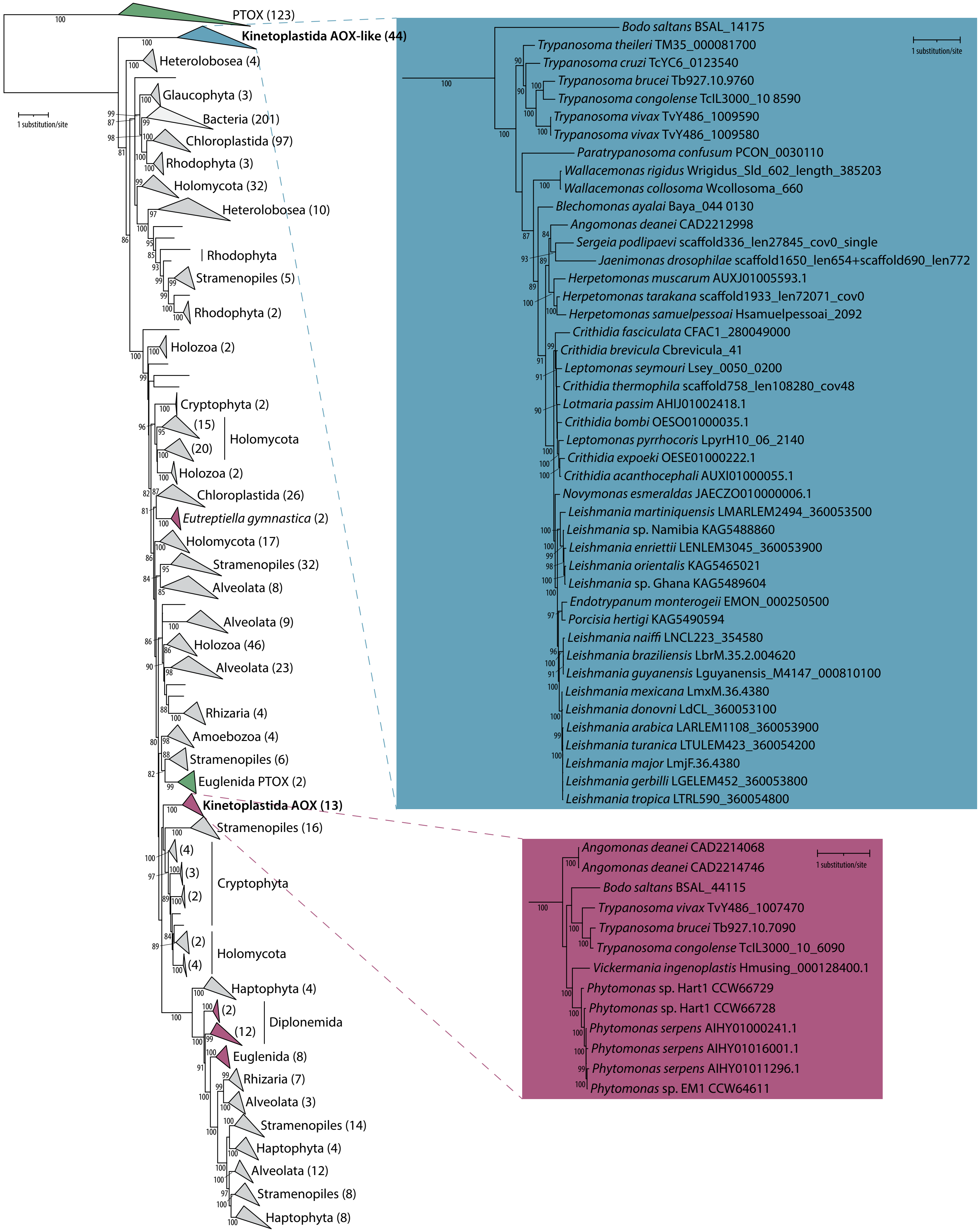

### Fig. S2

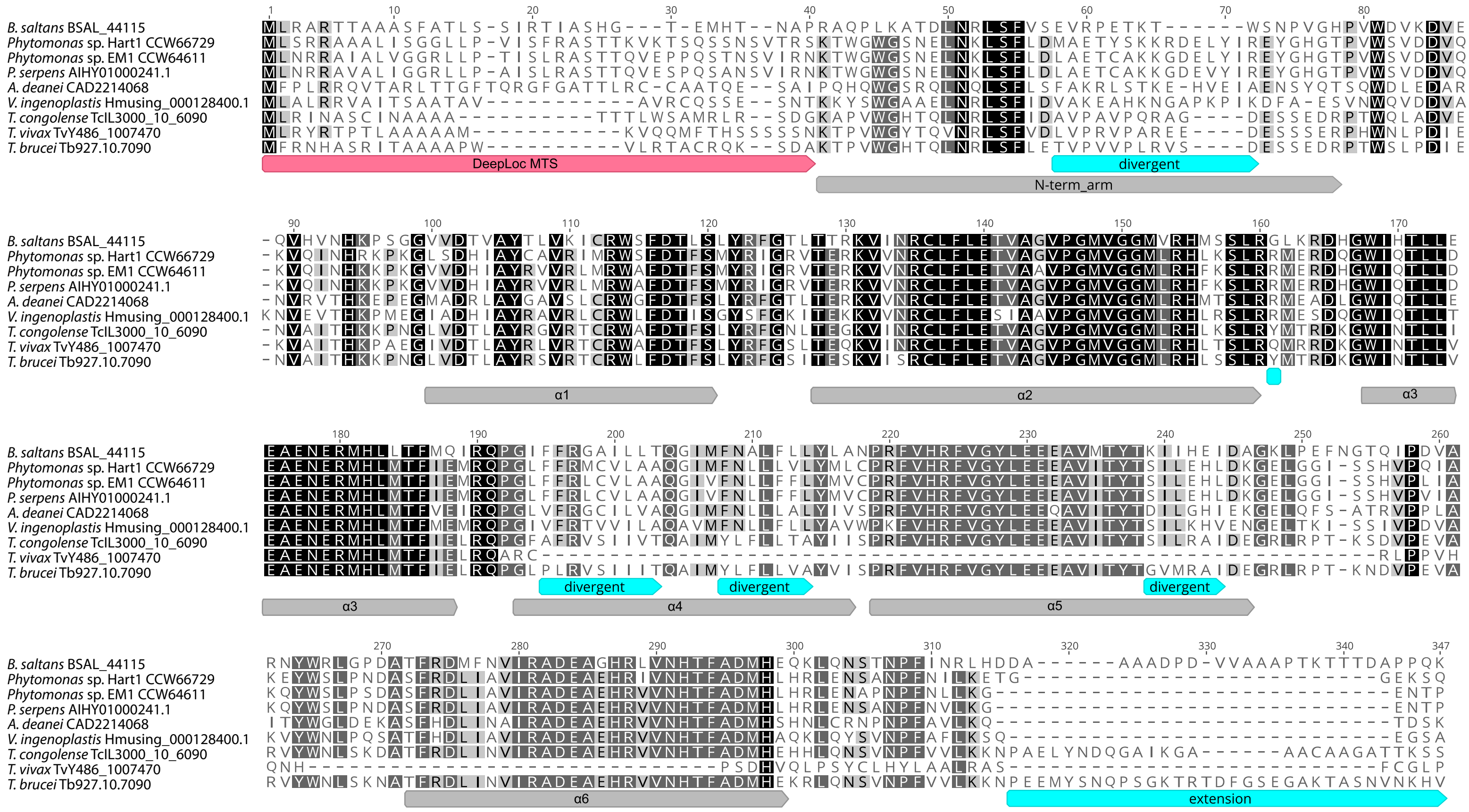
